# Detecting Long-term Balancing Selection using Allele Frequency Correlation

**DOI:** 10.1101/112870

**Authors:** Katherine M. Siewert, Benjamin F. Voight

## Abstract

Balancing selection occurs when multiple alleles are maintained in a population, which can result in their preservation over long evolutionary time periods. A characteristic signature of this long-term balancing selection is an excess number of intermediate frequency polymorphisms near the balanced variant. However, the expected distribution of allele frequencies at these loci has not been extensively detailed, and therefore existing summary statistic methods do not explicitly take it into account. Using simulations, we show that new mutations which arise in close proximity to a site targeted by balancing selection accumulate at frequencies nearly identical to that of the balanced allele. In order to scan the genome for balancing selection, we propose a new summary statistic, *β*, which detects these clusters of alleles at similar frequencies. Simulation studies show that compared to existing summary statistics, our measure has improved power to detect balancing selection, and is reasonably powered in non-equilibrium demographic models or when recombination or mutation rate varies. We compute *β* on 1000 Genomes Project data to identify loci potentially subjected to long-term balancing selection in humans. We report two balanced haplotypes - localized to the genes *WFS1* and *CADM2* - that are strongly linked to association signals for complex traits. Our approach is computationally efficient and applicable to species that lack appropriate outgroup sequences, allowing for well-powered analysis of selection in the wide variety of species for which population data are rapidly being generated.

## 1 Introduction

The availability of high-quality, population-level genomic data from a wide variety of species has spurred recent efforts to detect genomic regions subjected to natural selection (Vitti *et al.*, 2013, Xu *et al.*, 2015, Singh *et al.*, 2012). One type of pressure, balancing selection, occurs when more than one allele is maintained at a locus. This selection can arise from overdominance (in which the fitness of heterozygotes at a locus is higher than either type of homozygote) or from frequency-, temporally-, or spatially-dependent selection (Charlesworth, 2006). A classic case of overdominance occurs at the hemoglobin-*β* locus in populations located in malaria-endemic regions. Homozygotes for one allele have sickle-cell anemia, and homozygotes for the other allele have an increased risk of malaria. In contrast, heterozygotes are protected from malaria, and at most have a mild case of sickle-cell anemia (Aidoo *et al.*, 2002, Luzzatto, 2012).

The discovery of novel targets of balancing selection could help us better understand the role this selection has played in evolution, uncover traits that have been preserved for long evolutionary time periods, and aid in interpreting regions previously associated with phenotypes of interest. In addition, theory predicts that signatures of long-term balancing selection will be confined to regions of at most a few kilobases (Gao *et al.*, 2015). This feature of balancing selection’s targets leads to fewer possible causal variants than other types of selection, potentially aiding in understanding the underlying biology and associated mechanism.

Patterns of genetic variation around a locus targeted by balancing selection are distorted relative to a neutral locus. Because both alleles at a balanced locus are maintained in the population, the time to the most recent common ancestor (TMRCA) will be substantially increased if selection is maintained long enough (Charlesworth, 2006). This elevates the levels of polymorphism around the balanced locus and leads to a corresponding reduction in substitutions (*i.e.*, fixed differences relative to an outgroup species) (Charlesworth, 2006).

This deviation in the site frequency spectrum has been harnessed to identify signals of balancing selection in population data, genome-wide. These methods include Tajima’s D (Tajima, 1989), which detects an excess number of intermediate frequency alleles. Another commonly used method, the HKA test (Hudson *et al.*, 1987), uses the signal of high diversity and/or a deficit of substitutions. While these methods are easily implemented and widely applicable, their power under certain demographic scenarios or equilibrium frequencies is modest (DeGiorgio *et al.*, 2014).

If the selection began prior to the divergence of two species, then both species can share the balanced haplotype or variant (Charlesworth, 2006). These shared variants are unlikely to occur under neutrality (Gao *et al.*, 2015). Several recent studies have utilized primate outgroups to identify these trans-species polymorphisms (Andrés *et al.*, 2009, Leffler *et al.*, 2013, Teixeira *et al.*, 2015). While specific, this approach fails to identify selection if the balanced variant was lost in at least one of the species under consideration.

More powerful methods to detect balancing selection have been developed, though challenges have limited their broader application. DeGiorgio *et al.* (2014) proposed two model-based summaries, T1 and T2, which generate a composite likelihood of a site being under balancing selection. However, the most powerful measure (T2) requires the existence of a closely related outgroup sequence and knowledge of the underlying demographic)history from which an extensive grid of simulations must first be generated. New advances in estimating population-scale coalescent trees have also been harnessed to detect regions of the genome showing an unusually old TMRCA, but genome-wide application may be computationally prohibitive (Rasmussen *et al.*, 2014).

Despite these methodological advances, the exact frequencies of the excess intermediate frequency alleles seen under balancing selection have not been precisely quantified. The key insight motivating our work was the observation that the frequencies of these excess variants closely match the balanced allele’s frequency. We confirm this signature using simulations.

Motivated by this observation, and inspired by the structure of summary-spectrum based statistics (Tajima, 1989, Fay and Wu, 2000), we developed a new summary statistic that detects these clusters of variants at highly correlated allele frequencies. This statistic is computationally efficient and does not require knowledge of the ancestral state or an outgroup sequence. Using simulations, we show that our approach has equivalent or higher power to identify balancing selection than similar approaches, and retains power over a range of population genetic models and assumptions (i.e., demography, mutation, or recombination).

We report a genome-wide scan applying our statistic to humans using 1000 Genomes Project data (The 1000 Genomes Consortium, 2015), focusing on regions of high sequence quality. We find signals of balancing selection at two loci (*WFS1* and *CAMD2*) with functional evidence supporting these as the target genes, as well as signals at several previously known loci.

## 2 New Approaches

### 2.1 Allelic class build-up

We begin with an idealized model generating the expected distribution of allele frequencies around a balanced variant. Consider a new neutral mutation that arises within an outcrossing, diploid population. In a genomic region not experiencing selection, this mutation is expected to eventually either drift out of the population, or become fixed (*i.e.*, become a substitution). However, if the locus is under balancing selection, then the allele’s frequency can reach no higher than the frequency of the balanced allele it arose in linkage with, assuming no recombination (**Fig. 1**). This is because the frequency is constrained by selection. Without a recombination event and given enough time, variants that are fixed within these allelic classes (defined by the selected variant) accumulate (Charlesworth, 2006, Hey, 1991, Hudson, 1991). We used Wright-Fisher forward simulations to model neutral variants in a region closely linked to a variant under balancing selection (**Methods**). Within a region not expected to have experienced recombination since the start of the selection, we observed an excess number of variants with frequencies identical to that of the balanced variant, as predicted by this model (**Fig. 2a**).

**Figure 1:**
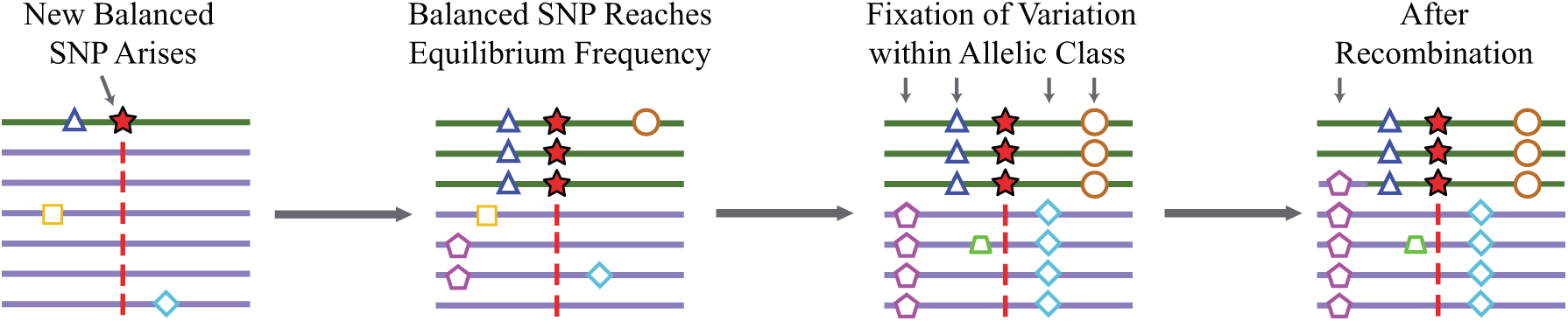
Model of allelic class build-up. (1) A new SNP (green star) arises in the population and is subject to balancing selection. (2) It sweeps up to its equilibrium frequency. (3) New SNPs enter the population linked to one of the two balanced alleles and some drift up in frequency. However, unlike in the neutral case, their maximum frequency is that of the balanced allele they are linked to, so variants build-up at this frequency (e.g. red diamond or brown circle). (4) Recombination decouples SNPs (e.g. pink pentagon) from the balanced site, allowing them to experience further genetic drift.

**Figure 2:**
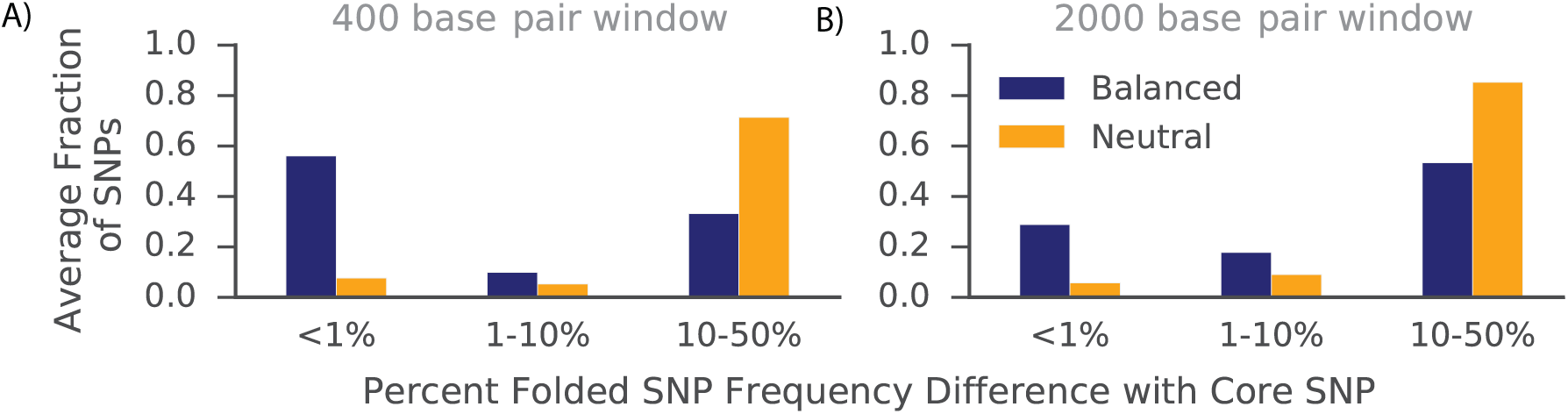
Simulations demonstrating build-up of alleles at frequencies similar to balanced alleles as compared to non-balanced counterparts. The blue bars indicate the fraction of SNPs in simulation replicates at specific frequency differences away from a balanced core site. In contrast, the orange bars represent simulation replicates that lack a balanced variant. Instead, the core site is chosen to be a neutral variant within frequency 10% of the equilibrium frequency of variants introduced in the balanced simulations (A) Folded frequency differences between the core SNP and each other SNP in a 400 base-pair window surrounding the core site. Recombination is not expected to have occurred in this region since the start of selection (Gao *et al.*, 2015). (B) Frequency differences in 2, 000 base-pair windows, where recombination is expected to have occurred since the start of selection.

Eventually, recombination decouples variants from the balanced allele, which allows them to drift to loss or fixation within the population. However, even after recombination, the frequency of the variants previously fixed in their allelic class will remain close to that of their previous class until enough time has passed for genetic drift to significantly change their frequencies (Charlesworth, 2006, Hey, 1991, Hudson, 1991). In our simulations of balancing selection, a window expected to have experienced recombination since selection’s start still has an excess number of variants at similar frequencies to the balanced variant. However, there is a smaller excess at identical frequencies relative to the more narrow window, demonstrating the effects of recombination (**Fig. 2b**).

### 2.2 A Measure for Allele Frequency Correlation

To capture this signature, we derive a measurement of frequency similarity between a core variant and a second variant of interest. Let *n* be the number of chromosomes sampled, *f*_0_ be frequency of the core SNP, *f_i_* be the frequency of the second SNP, *i*, and *p* be a scaling constant (see **Supplementary note**). Finally, *g*(*f*) returns the folded allele frequency and *m* is the maximum possible folded allele frequency difference between the core SNP and SNP *i*, We then measure the similarity in frequency, *d_i_*, by:

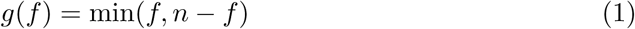

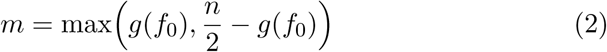

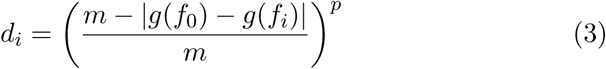

Thus, *g*(*f*_0_) − *g*(*f_i_*) is the folded frequency difference between the core SNP and the SNP under consideration. We then subtract this value from *m*, the maximum folded frequency difference possible with the core SNP, and then divide by *m*. This gives the fraction of the maximum folded frequency difference of the SNP under consideration compared with the core SNP. We then raise it to the power *p* so that we can weight variants in a non-linear fashion with respect to this fraction. Therefore, *d_i_* can range from 0 if a SNP has the maximum frequency difference with the core SNP, to 1 if SNP *i* is at the same frequency as the core SNP. We give guidance on the choice of *p* in the Online Supplement. However, the power of *β* is fairly insensitive to its value (Fig. S12). We use the folded site frequency spectrum in calculating *d_i_*, as the frequency difference between the core variant and the second variant is independent of whether the derived or ancestral allele of the nearby allele is in linkage with the derived or ancestral core allele.

In a region under long-term balancing selection, the average *d_i_* between a core SNP and the surrounding variants is expected to be elevated. However, *d_i_* alone is not optimally powered to detect balancing selection, as its value will be sensitive to changes in the mutation rate in the surrounding region, and it does not take into account the probability of seeing each allele frequency under neutrality.

### 2.3 Capturing Allelic Class Build-up

We propose a statistic, *β*, that uses our measure of allele frequency correlation, *d_i_*, combined with a measure of the overall mutation rate, to detect balancing selection. Our approach is inspired by previous summary statistics of the site frequency spectrum (Tajima, 1989, Fay and Wu, 2000). These methods compute the difference between two estimators of *θ*, the population mutation rate parameter, one of which is more sensitive to characteristics of the site frequency spectrum distorted in the presence of natural selection. We propose to calculate *β* at each SNP in a region of interest to identify loci in which there is an excess of variants near the core SNP’s allele frequency, as evidence of balancing selection.

It has been previously shown that the mutation rate in a region can be estimated as: 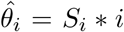 where *S_i_* is the total number of derived variants found *i* times in the window from a sample of *n* chromosomes in the population (Fu, 1995). An estimator of *θ* can then be obtained by taking a weighted average of *θ_i_*. In our method, we weight by the similarity in allele frequency to the core SNP, as measured by *d_i_*. If there is an excess of variants at frequencies close to the core SNP allele frequency, then our new estimator, *θ_β_*, will be elevated. We propose:

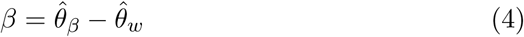

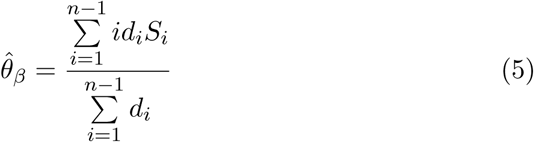

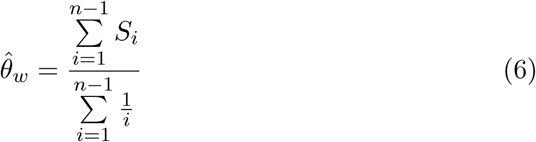

*θ_ω_* is simply Watterson’s estimator (Watterson, 1975 *β* is, in effect, a weighted average of SNP counts based on their frequency similarity to the core SNP. We exclude the core site from our estimation of *θ* _*ω*_ and *θ_β_*

To better understand the properties of *β*, we used simulations to examine its distribution with and without a balanced SNP (**Fig. S2**). As expected, under long-term balancing selection *β* tends to be greater than zero, and under neutrality it tends to be close to zero.

We note that the mean value of *β* in our neutral simulations generally increases slightly with higher equilibrium frequencies. This behavior is expected because higher frequency alleles will tend to have a longer TMRCA and therefore higher diversity. The exception to this trend is neutral SNPs of frequency 0.5, which we posit is due to the fact that this allele frequency requires the most time for mutations to drift up to the equilibrium frequency needed to fix in their allelic class.

In this version of *β*, knowledge of the ancestral state for each variant is required. To address this possible shortcoming, we developed a version of the statistic based on a folded site frequency spectrum. This formulation is available in the **Online Supplement**.

While our statistic can be calculated on any window size, previous work has suggested that the effects of balancing selection localize to a narrow region surrounding the balanced site (Gao *et al.*, 2015). Ultimately, the optimal window size depends on the recombination rate, as it breaks up allelic classes. In the **Online Supplement**, we present some mathematical formulations to suggest reasonable window sizes.

## 3 Results

### 3.1 Power Analysis

We used forward simulations (Haller and Messer, 2017) to calculate the power of our approach to detect balancing selection relative to other commonly utilized statistics. Initially, we simulated a single, overdominant mutation for each simulation replicate in an equilibrium demographic model, varied over a range of balancing selection equilibrium frequencies and onset times (**Methods**). We also simulated genomic regions in which all variants were selectively neutral. We then computed the power of *β*, Tajima’s D, HKA, and T1 to distinguish between simulation replicates with a balanced variant (our balanced simulations) or those with only neutral mutations (our neutral simulations). As a reference, we also measured the likelihood-based statistic, T2.

*β*, Tajima’s D and HKA use a sliding window approach, in units of base pairs, when scanning the genome, while T1 and T2 use the number of informative sites (polymorphisms plus substitutions). In order to make a fair comparison between these methods, we first determined the most powerful window size for each method using simulations (**Fig. S5**). For the summary statistics, a 1kb window size did well across a range of selection timings and equilibrium frequencies. This 1kb region matches the approximate size of the ancestral region, in which there have been no expected recombination events between allelic classes (**Supplementary Information**).

For T1 and T2, a number of informative sites of approximately 20, or 10 on either side of the core site, achieved maximum power in simulations (**Fig. S6**). Furthermore, this roughly matches the expected number of informative sites in a 1kb region under selection (**Supplementary Information**). Therefore, a window of 20 total informative sites is roughly equal to the expected ancestral region size, which is roughly equal to the window at which all these methods achieve optimal power. For this reason, we used a 1kb window or 20 informative sites, as applicable, when calculating each statistic.

Compared to other summaries, *β* had the greatest performance across most parameter combinations (**Fig. 3, Fig. S4-17**). As expected, *β* performs slightly worse than T2 under many conditions. However, unlike T2, our method does not require an outgroup sequence, or grids of simulations which are computationally expensive.

**Figure 3:**
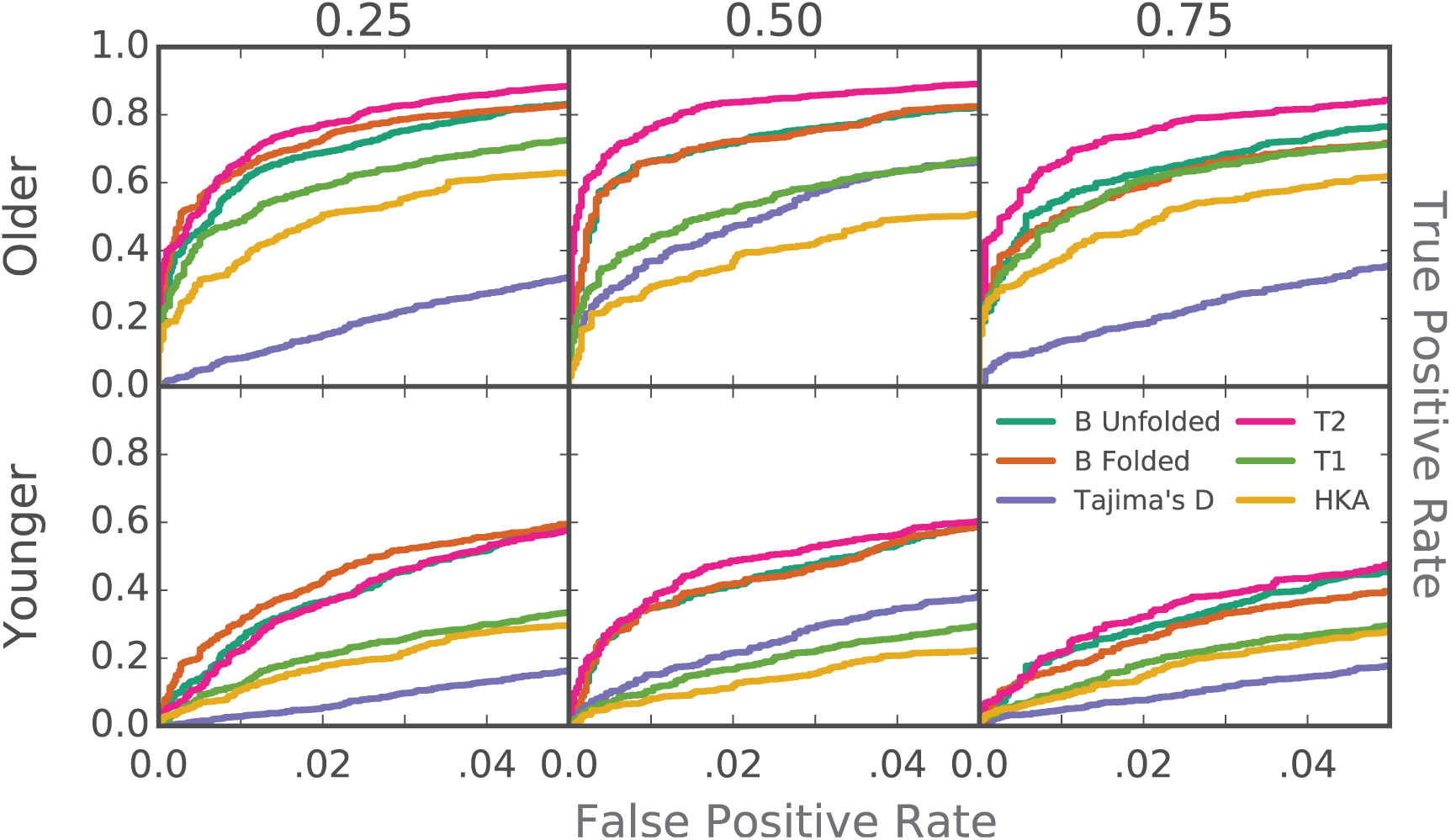
Power of methods to detect ancient balancing selection. Power was calculated based on simulation replications containing only neutral variants (True Negatives) or containing a balanced variant that was introduced (True Positives). Columns correspond to simulations of balanced alleles at equilibrium frequencies .25, .50 and .75. Rows correspond to older and more recent selection, beginning 250,000 and 100,000 generations prior to sampling, respectively.

We next investigated the power of *β* under more complex demographic scenarios (Methods) compatible with recent human history (DeGiorgio *et al.*, 2014). We found that *β* performs well under bottleneck and expansion models. Under an expansion scenario, the performance of all methods decreased (**Fig. S7**), consistent with results from previous studies (DeGiorgio *et al.*, 2014), possibly due to the larger population size increasing the expected time until an allele can fix in its allelic class. The effect of a population bottleneck on power was less drastic and led to a slight increase in power to detect more recent selection (**Fig. S8**).

Population substructure can confound scans for selection (Ingvarsson, 2004, Schierup *et al.*, 2000). To investigate the power of our method in these scenarios, we simulated two models of population substructure. First, we considered a model of two completely subdivided populations. We pooled together 50 individuals from each subpopulation with which to perform the statistical calculations. In this case, the power of all methods to detect balancing selection at equilibrium frequency 0.5 decreased considerably (**Fig. S9**). This matches expectation, as this situation is expected to drastically increase the number of variants at frequency 0.5.

Next, we considered a two-pulse model of ancient admixture. We selected this model because of its approximation of Neanderthal admixture into human (Vernot and Akey, 2015), which may be thought to confound scans for selection in humans. Power with Neanderthal admixture stayed roughly the same as without (**Fig. S10**). This is as expected, as most haplotypes introduced through admixture are expected to be at very low frequency.

We next examined the power for all methods under models of variable mutation rates, recombination rates, and sample sizes. As expected, the power of all methods was positively correlated with mutation rate (**Fig. S13 and S15**), and negatively correlated with recombination rate (**Fig. S14 and S16**). A higher mutation rate provides more variants that can accumulate within an allelic class, whereas a lower recombination rate allows for longer haplotypes upon which mutations can accumulate.

*β* has reasonable power down to very small sample sizes, achieving near maximum power with as few as 20 sampled chromosomes (**Fig. S19 and S20**). In practice, the sample size used to calculate the frequency of each variant may differ between variants. We tested the power of *β* when the sample size of each variant is downsampled from the original size of 100 by a random amount from 0 to 25 individuals. We found that this decreases power very slightly, and that lower values of *p* perform better in this scenario (**Fig. S11**).

Finally, power remained high under frequency-dependent selection (**Fig S18**), and when a lower selection coefficient was simulated (**Fig. S17**). This matches expectation, as frequency-dependent selection is expected to maintain haplotypes in the population for long time periods, causing allelic class build-up. A lower selective coefficient would be expected to lower the probability of maintenance of the balanced allele in the population, but conditioned on this maintenance, should not affect power, as we observed.

Simulations show that the power of the folded version of *β* is similar to the unfolded version at intermediate allele frequencies, but has reduced power at very high frequencies (**Fig. S4**). However, even at these frequencies, it still outperforms Tajima’s D, the only other statistic of those tested which does not require knowledge of the ancestral state or an outgroup.

### Genome-wide Scan in Human Populations

We applied the unfolded version of *β* to population data obtained by the 1000 Genomes Project (Phase 3) to detect signatures of balancing selection (The 1000 Genomes Consortium, 2015). We calculated the value of *β* in 1kb windows around each SNP in all 26 populations, separately. We focused on regions that passed sequencing accessibility and repeat filters (**Methods**).

In addition, we filtered out variants which did not have a folded frequency of at least 15% in a minimum of one population. The purpose of the frequency filter is to prevent false positives: we were unable to simulate balancing selection with a folded equilibrium frequency of less than 15%, due to the high frequency of one allele drifting out of the population. Although this phenomenon has not been described for population sizes near that of humans to our knowledge, it has been detailed for lower effective population sizes (Ewens and Thomson, 1970). Therefore, it seems unlikely that a balanced variant with a folded equilibrium frequency less than 15% could successfully be maintained in a population.

We defined *β* extreme scores as those in the top 1% in the population under consideration (**Methods**). We analyzed the autosomes and Xchromosome separately. Because our method is substantially better powered to detect older selection, we focus on signals of selection that predate the split of modern populations. For this reason, we further filtered for loci that were top-scoring in at least half of the populations tested (**Methods**). We focus on results of our *β* unfolded scan, however, we also scanned using the folded *β* statistic to test for robustness of our top scoring sites.

We identified 8,702 autosomal, and 317 X-chromosomal, top-scoring variants that were shared among at least half (≥ 13) of the 1000 Genomes populations (**Supplementary Data**). Together, these variants comprise 2,453 distinct autosomal and 86 X-chromosomal loci, and these signatures overlapped 692 autosomal and 29 X-chromosomal genes.

### 3.2 Characterization of Identified Signals

Trans-species haplotypes are two or more variants that are found in tight linkage and are shared between humans and a primate outgroup (in our case, chimpanzee). These haplotypes are highly unlikely to occur by chance, unlike trans-species SNPs, which are expected to be observed in the genome due to recurrent mutations (Gao *et al.*, 2015). These haplotypes present a signature of balancing selection independent from the signature captured by *β*. If *β* captures true signatures of balancing selection, one would expect an enrichment of high values at trans-species haplotypes. We found that *β* was in fact predictive of trans-species haplotype status from Leffler *et al.* (2013), even after including adjustments for the distance to the nearest gene (*P <* 2 × 10 ^*-16*^, **Methods**).

Our scan identified several loci that have been previously implicated as putative targets of balancing selection (**Supplementary Data**). Several major signals occurred on chromosome 6 near the HLA, a region long presumed to be subjected to balancing selection (Hedrick, 1998, Hughes and Nei, 1988). In particular, we found a strong signal in the HLA at a locus influencing response to Hepatitis B infection, rs3077 (Jiang *et al.*, 2015, De-Giorgio *et al.*, 2014, Thursz *et al.*, 1997). Several additional top sites in our scan matched those from DeGiorgio *et al.* (2014). These include sites that tag phenotypic associations (Welter *et al.*, 2014), such as *MYRIP*, involved with sleep-related phenotypes (Gottlieb *et al.*, 2007), and *BICC1*, associated with corneal astigmatism (Lopes *et al.*, 2013). We focus on two of our top-scoring regions, located in the *CADM2* and *WFS1* genes. In addition to passing the 1000 Genomes strict filter and the RepeatMasker test, these haplotypes also passed Hardy-Weinberg filtering (**Methods**).

### 3.3 A signature of selection at the *CADM2* locus

One of our top-scoring regions fell within an intron of the cell adhesion molecule 2 gene, *CADM2*. This locus contains a haplotype with *β* scores falling in the top 0.25 percentile in 17 of the 1000 Genomes populations, and scoring in the top 0.75 percentile across all 26 populations (Fig. 4). This site was also a top scoring SNP in the CEU population based on the T2 statistic (DeGiorgio *et al.*, 2014). In our scan using the folded *β* statistic, this haplotype contained top-scoring variants in 20 populations, indicating the result was not due to ancestral allele miscalling. In the remaining six populations, the haplotype was at folded frequency 0.15 or lower, where the folded version of *β* has significantly reduced power.

**Figure 4:**
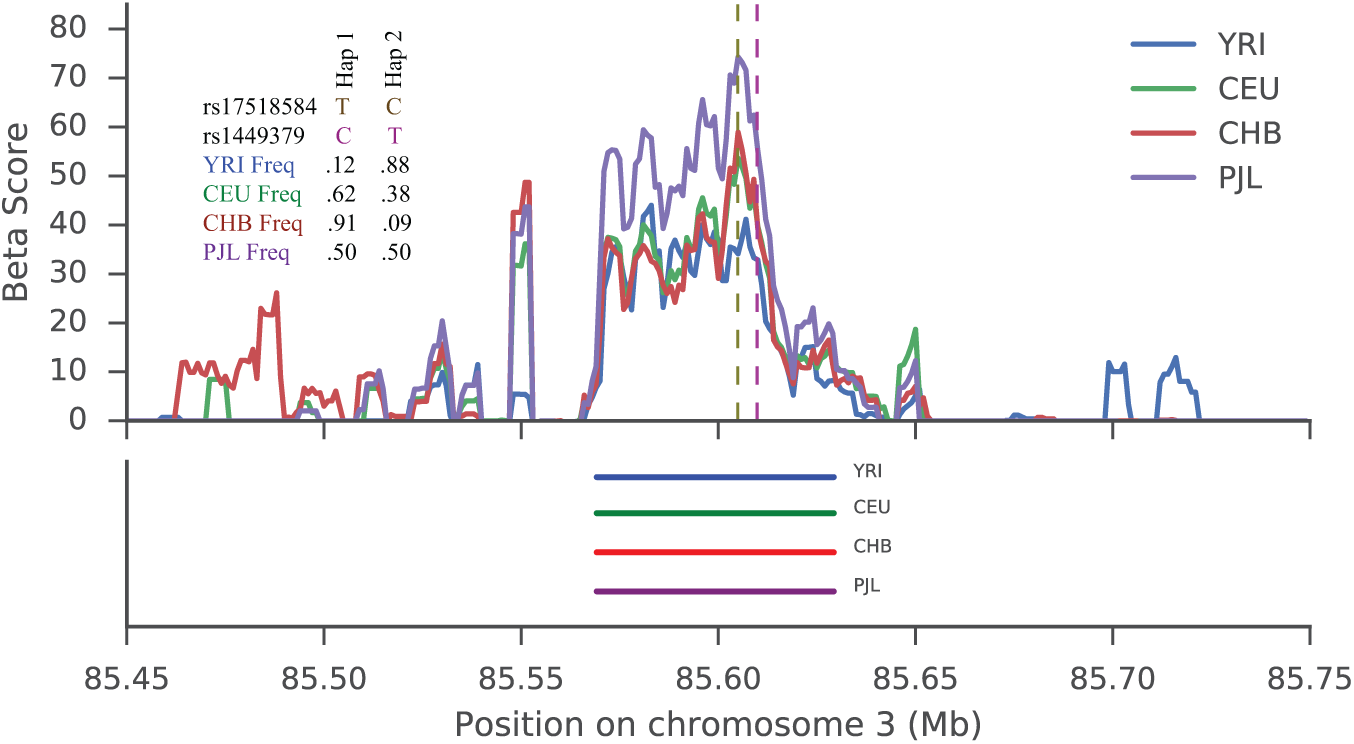
Signal of balancing selection at *CADM2*. The signal of selection is located in an intron of *CADM2*. (a) rs17518584 is the lead GWAS SNP for several intellectual traits and is marked by the brown vertical dashed line. The purple dashed line marks two regulatory variants found on the balanced haplotype. *β* scores were calculated using a rolling average with windows of size 5 kb, including only SNPs at the same frequency as the core SNP in the average. In addition, we show the allele frequencies of the GWAS and a top-scoring *β* SNP in each representative population. (b) Approximate haplotype lengths for each population.

To elucidate the potential mechanisms contributing to the signal in this region, we overlapped multiple genomic datasets to identify potential functional variants that were tightly linked with our haplotype signature. First, one variant that perfectly tags (EUR *r*^2^ = 1.0) our signature, rs17518584, has been genome-wide significantly associated with cognitive functions, including information processing speed (Davies *et al.*, 2015, Ibrahim-Verbaas *et al.*, 2016). Second, multiple variants in this region co-localized (EUR *r*^2^ between 0.9 *-* 1 with rs17518584) with eQTLs of *CADM2* in numerous tissues (Lung, Adipose, Skeletal Muscle, Heart-Left Ventricle), though notably not in brain (The GTEx Consortium, 2015). That said, several SNPs with regulatory potential (RegulomeDB scores of 3a or higher) are also strongly tagged by our high-scoring haplotype (EUR *r*^2^ between 0.9 *-* 1.0 with rs17518584), which include regions of open chromatin in Cerebellum and other cell types (Boyle *et al.*, 2012). Several SNPs on this haplotype, particularly rs1449378 and rs1449379, fall in enhancers in several brain tissues, including the hippocampus (Ernst and Kellis, 2012, Boyle *et al.*, 2012). Taken collectively, these data suggest that our haplotype tags a region of regulatory potential that may influence the expression of *CADM2*, and potentially implicates cognitive or neuronal phenotypes in the selective pressure at this site.

### 3.4 A signature of balancing selection near the diabetes associated locus, *WFS1*

We identified a novel region of interest within the intron of *WFS1*, a trans-membrane glycoprotein localized primarily to the endoplasmic reticulum (ER). *WFS1* functions in protein assembly (Takei *et al.*, 2006) and is an important regulator of the unfolded protein and ER Stress Response pathways (Fonseca *et al.*, 2005). A haplotype in this region (approximately 3.5 kb) contains approximately 26 variants, 3 of which are in high-quality windows and are high-scoring *β* in all populations (Fig. 5). The haplotype was also in the top 1 percentile in our folded *β* scan in 21 populations. In the remaining 5 populations, this haplotype was at frequency 0.82 or higher, where the folded version of has significantly lower power than the unfolded version.

**Figure 5:**
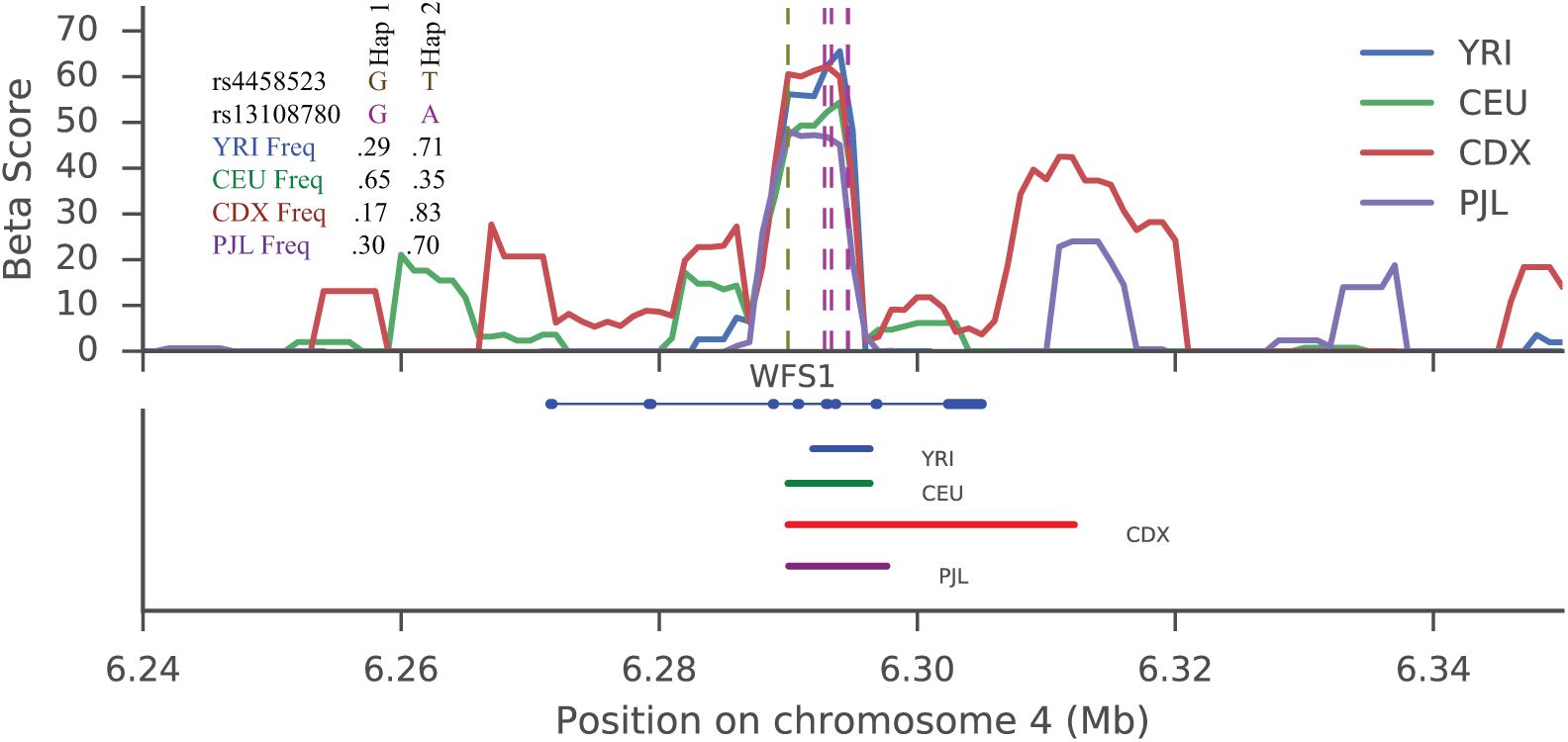
Signal of balancing selection at the WFS1 gene. (a) rs4458523 is the lead GWAS SNP for diabetes, and is marked by the brown vertical dashed line. The purple dashed line marks 5 regulatory variants found on the balanced haplotype. In addition, we show the allele frequencies of the GWAS and a top-scoring *β* SNP in each representative population. (b) Approximate haplotype lengths for each population.

Our identified high-scoring haplotype tags several functional and phenotypic variant associations. First, one variant that perfectly tags our signature (EUR *r*^2^ = 1.0), rs4458523, has been previously associated with type 2 diabetes (Voight *et al.*, 2010, Mahajan *et al.*, 2014). Second, multiple variants in this region are associated with expression-level changes of *WFS1* in numerous tissues (The GTEx Consortium, 2015); these variants are strongly tagged by our high-scoring haplotype (EUR *r*^2^ between 0.85 *-* 0.9 with rs4458523). Finally, several SNPs with regulatory potential (RegulomeDB scores of 2b or higher) are also strongly tagged by our high-scoring haplotype (EUR *r*^2^ between 0.9 *-* 1.0 with rs4458523). Taken collectively, these data suggest that our haplotype tags a region of strong regulatory potential that is likely to influence the expression of *WFS1*.

## 4 Discussion

Informed by previous theory on allelic-class build-up (Charlesworth, 2006, Hey, 1991, Hudson, 1991), we developed a novel summary statistic to detect the signature of balancing selection, and measured efficacy and robustness of our approach using simulations. While our method does not require knowledge of ancestral states for each variant from outgroup sequences, this information can improve power at extreme equilibrium frequencies.

Although our method outperforms existing summary statistic methods, it is not as powerful as the computationally-intensive approach of T2, which uses simulations to calculate likelihoods of observed data (DeGiorgio *et al.*, 2014). To improve power, we considered utilizing information on rates of substitutions, but this did not substantially improve discriminatory power (**Online Methods**). Alternative possibilities could include (i) consideration of the region past the ancestral region surrounding the balanced variant, or (ii) deviations in the frequency spectrum beyond just near identical frequencies to the balanced SNP. As expected from theory, we also note that models of population structure can also produce our haplotype signature, emphasizing the requirement to perform scans on individual populations.

Balancing selection can cause a similar signature in self-fertilizing species, though we focused on out-crossed species in this report. Previous work has shown that given the same selection coefficient, the signature of balancing selection can be wider in self-fertilizing species due to a lower effective recombination rate (Nordborg *et al.*, 1996). However, lower recombination rate also means that background selection leaves a wider footprint on the genome in these species, which can reduce levels of polymorphism (Agrawal and Hartfield, 2016). Furthermore, a decrease in the frequency of heterozygotes, owing to selfing, can reduce or eliminate the effects of heterozygote advantage. Instead, modes of balancing selection like frequency, temporally or spatially-dependent selection may be more significant.

We have also assumed a single causal variant throughout. However, there may be more than one variant at a locus experiencing balancing selection. This situation is thought to occur throughout the HLA region (Hedrick, 1998). Assuming the maintenance of multiple variants, this scenario would also increase the regional TMRCA, leading to allele class build-up, spanning perhaps a larger window than our single-variant models (Lenz *et al.*, 2016). The dynamics of this type of situations could be the focus of future work.

Although it is impossible to know the true selective pressure underlying our highlighted loci, our results suggest that balancing selection could contribute to the genetic architecture of complex traits in human populations. At the *CADM2* locus, functional genomics data suggests that our haplotype signature may connect to brain-related biology. Intriguingly, a recent report also noted a strong signature of selection at this locus in canine (Freedman *et al.*, 2016), suggesting a possibility of convergent evolution. That said, the phenotypes that have resulted in a historical fitness trade-off at this locus are far from obvious.

Similarly, speculation on the potential phenotypes subject to balancing selection at *WFS1* should also be interpreted cautiously. It is known that autosomal recessive, loss of function mutations in this gene cause Wolfram Syndrome. This gene is a component of the unfolded protein response (Fonseca *et al.*, 2005) and is involved with ER maintenance in pancreatic β-cells. Furthermore, deficiency of *WFS1* results in increased ER stress, impairment of cell cycle, and ultimately increased apoptosis of beta-cells (Yamada *et al.*, 2006). These data would suggest that reduced expression of *WFS1* would be diabetes risk increasing; however, eQTLs that co-localized with the diabetes risk-increasing allele *elevate* expression, at least in non-pancreas tissue, suggesting perhaps a more complex functional mechanism. Furthermore, how the unfolded protein response could connect to historical balancing selection is also not immediately obvious. One possibility derives from recent work suggesting that these pathways respond not only to stimulus from nutrients or ER stress, but also to pathogens (Nakamura *et al.*, 2010). This could suggest the possibility that expression of *WFS1* is optimized in part to respond to pathogen exposure at a population level.

*β* is powered to detect balancing selection when outgroup sequences are not available and can do so quickly and easily. Given the increasing ease of collecting population genetic data from non-model organisms, our approach is in a unique position to characterize balancing selection in these populations.

An implementation of both the folded and unfolded versions of *β* is available for download at <u>https://github.com/ksiewert/BetaScan</u>.

## 5 Materials and Methods

### 5.1 Simulations

Simulations were performed using the forward genetic simulation software SLiM 2.0 (Haller and Messer, 2017). In our simulations, neutral mutations and recombination events occur at a predefined rate throughout the entire length of the simulation. A burn-in time of 100,000 generations was first simulated to achieve equilibrium levels of variation. Then, two populations representing humans and chimpanzees split from this original population, and were simulated for 250,000 additional generations. We then sampled 100 chromosomes from the human population, and 1 chromosome from the chimpanzee population. We first simulated these scenarios under parameters suitable for human populations, with mutation and recombination rates of π = *r* = 2.5 × 10 ^-8^ and *N_e_* = 10, 000.

We generated two sets of simulations: one without a balanced variant (the set we refer to as our neutral simulations) and one with a balanced variant (balanced simulations). In the second set, a single balanced variant was introduced at the center of the simulated region in the human population, either at the time of speciation (250,000 generations prior to simulation ending), or 150,000 generations after speciation (100,000 generations prior to simulation ending). The simulations then continued as normal, conditional on maintenance of the balanced SNP in the population. If this balanced variant was lost, the simulation restarted at the generation in which the balanced variant was introduced. In the second (neutral) set, no balanced variant was introduced, so all variants are selectively neutral.

Each balanced SNP had an overdominance coefficient *h* and selection coefficient *s*. The fitness of the heterozygote is then 1 + *hs*, and the fitness of the ancestral and derived homozygotes are 1 and 1 + *s*, respectively. We simulated two different *s* values: 10 ^-2^ (our default) and 10 ^-4^. We simulated six different equilibrium frequencies: 0.17, 0.25, 0.5, 0.75, 0.83, which correspond to *h* = *-*0.25, *-*0.5, 100, 1.5, 1.25. Negative *h* values were paired with negative *s* values.

After simulation completion, the frequency of each variant in the sampled individuals was calculated. Substitutions were defined as any variant in which the allele from the chimpanzee chromosome was not found in the sampled human individuals. For each set of balanced simulations, we define the core SNP as the variant under balancing selection. For each set of balanced simulations, we then find a corresponding set of core SNPs in our neutral simulations which are within frequency 10% of the equilibrium frequency of the balanced variants. We then calculated the score for each statistic on these core variants. In this way, we have statistic scores for the balanced variant from each balanced simulation replicate, and a score for a neutral variant matched for similar frequency. For more details on how each statistic was calculated, see (**Supplementary Methods**).

To increase simulation speed, we rescaled our simulations by a factor of 10 for specified power analyses in the supporting information (Hoggart *et al.*, 2007); results presented in the main text were not rescaled. A minimum of 1500 simulation replicates were performed for each parameter set. We simulated 10kb regions for each simulation replicate, with the exception of the analysis of optimal windows size, in which case a 100kb region was simulated.

### 5.2 Empirical Site Analysis

To apply our method to 1000 Genomes data, we first downloaded data for each of the 26 populations in phase 3 of the project (obtained May 2nd, 2013). We then calculated allele frequencies separately for each population, and calculated *β* in 1 kb sized windows centered around each SNP for each population.

Because poorly sequenced regions can artificially inflate the number of SNPs in a region, we then filtered out regions that contained one or more base pairs that were ruled as poor quality in the 1000 Genomes phase 3 strict mask file. For further confirmation that the signal was not a result of poor mapping quality, we overlapped SNPs of interest with hg19 human Repeat-Masker regions, downloaded from the UCSC Table Browser on February 9th, 2017. We then removed all core SNPs from consideration that were found within a repeat, similar to Bubb *et al.* (2006). We further removed SNPs that were not of common frequency (at or above a folded frequency of 15%) in at least one population. After filtering, there were 1,803,299 SNPs that remained. We then found the top 1% of these high-quality SNPs in each population in our *β* scan.

Unknown paralogs or other technical artifacts could inflate the number of intermediate frequency alleles. Although the 1000 Genomes data provides strict quality filter masks, we wanted to further verify that our haplotypes of interest in *WFS1* and *CADM2* were not the result of obvious technical artifacts. In order to do this, we used the −−hardy fiag in vcftools (Danecek *et al.*, 2011), and investigated both the one-tailed p-value for an excess of heterozygotes, and the two-tailed p-value, in our 4 representative populations (YRI, CEU, CDX and PJL). All variants on these haplotypes had p-values above 1 *×* 10^−3^.

The lowest autosomal significance cut-off of any population, ASW, corresponds to a *β* score of 47.49. This score is in the top 0.05 percentile of core SNPs in neutral simulations corresponding to an equilibrium frequency of 0.5 (**Fig. S3**).

To find top-scoring sites that are also GWAS hits, we obtained LD proxies in European populations for our top-scoring SNPs, using a cut-off of *r*^2^ of 0.9, a maximum distance of 50kb and a minimum minor allele frequency of 5%. We then overlapped these LD proxies with GWAS hits obtained from the GWAS Catalog to get our final list of putatively balanced GWAS hits (Welter *et al.*, 2014). Gene names and locations were downloaded from Ensembl BioMart on November 26th, 2016.

For our trSNP comparison, we used the Human/Chimp shared haplotypes from Leffler *et al.* (2013). Using logistic regression, we then modeled the outcome of a SNP being part of a trHap as dependent on the *β* Score and distance to nearest gene.

## 6 Supplementary Material

Supplementary figures S1-S20, and Supplementary methods are available on bioRxiv.

## 7 Acknowledgments

We would like to acknowledge the members of the Voight Lab for their helpful suggestions and support, Philipp Messer for his comments on this manuscript, and comments from two anonymous reviewers. This work was supported through grants from the National Institutes of Health (NIDDK R01DK101478 to BFV and T32HG000046-17 to KMS) and the Alfred P. Sloan Foundation (BR2012-087 to BFV).

